# Asymptomatic gut colonization by extended-spectrum beta-lactamase-producing *Escherichia coli* is not associated with an altered gut microbiome or metabolome in Dutch adults

**DOI:** 10.1101/2021.05.18.444613

**Authors:** Q. R. Ducarmon, R. D. Zwittink, R. P. J. Willems, A. Verhoeven, S. Nooij, F.R.M. van der Klis, E. Franz, J. Kool, M. Giera, C. M. J. E. Vandenbroucke-Grauls, S. Fuentes, E. J. Kuijper

## Abstract

**Background:** Gut colonization by antibiotic resistant *E. coli* strains, including extended-spectrum beta-lactamase (ESBL)-producing *E. coli* is a risk factor for developing overt infection. The gut microbiome can provide colonization resistance against enteropathogens, but it remains unclear whether it confers resistance against potentially pathogenic ESBL-producing *E. coli*.

**Materials:** From a Dutch cross-sectional population study (PIENTER-3), feces from 2751 individuals were used to culture ESBL-producing bacteria. Of these, we selected 49 samples which were positive for an ESBL-producing *Escherichia coli* (ESBL^+^), and negative for a variety of variables known to affect microbiome composition. These were matched in a 1:1 ratio to ESBL^−^ samples based on age, sex, having been abroad in the past six months and ethnicity. Shotgun metagenomic sequencing was performed and taxonomic species composition and functional annotations (microbial metabolism and carbohydrate-active enzymes) were determined. Targeted quantitative metabolic profiling (^1^H NMR-spectroscopy) was performed to investigate metabolomic profiles.

**Results:** No differences in alpha or beta diversity were observed, nor in relative abundance, between ESBL^+^ and ESBL^−^ individuals based on bacterial species level composition. Machine learning approaches based on microbiota composition did not accurately predict ESBL status (area under the receiver operating characteristic curve (AUROC)=0.53), neither when based on functional profiles. The metabolome did also not convincingly differ between ESBL groups as assessed by a variety of approaches, including machine learning through random forest (AUROC=0.61).

**Conclusion:** Using a combination of multi-omics and machine learning approaches, we conclude that asymptomatic gut carriage of ESBL-producing *E. coli* is not associated with an altered microbiome composition or function. This may suggest that microbiome-mediated colonization resistance against ESBL-producing *E. coli* is not as relevant as it is against other enteropathogens.

## INTRODUCTION

*Escherichia coli* is a common gut commensal, but several strains possess virulence factors that enable them to cause gastrointestinal, urinary and extraintestinal infections^1, 2^. Colonization of the gut by multidrug-resistant organisms (MDRO), including extended-spectrum beta-lactamase (ESBL)-producing *E. coli* and carbapenem-resistant *E. coli*, often precede infections^3^. The gut microbiome can mediate colonization resistance against several enteric pathogens, but it remains unclear whether this is also the case for MDROs such as ESBL-producing *E. coli*, especially since many individuals harbor commensal *E. coli*. Colonization resistance can be conferred by the gut microbiome through nutrient competition, production of antimicrobial compounds, support of gut barrier integrity, bacteriophage deployment and through interaction with the immune system^4^. However, studies in humans have reported conflicting evidence regarding which bacterial genera or species within the gut microbiome could be of relevance in providing colonization resistance against ESBL-producing *E. coli* or ESBL-producing *Enterobacterales*. These conflicting results can, at least partially, be traced back to several confounding factors (e.g. medication) in those studies^5–8^. It was recently shown that unevenly matched case-controls studies with regard to lifestyle and physiological characteristics can produce spurious microbial associations with human phenotypes like disease, or in this case, colonization by ESBL-producing *E. coli*^9^.

Here, we aimed to compare the gut microbiome and metabolome between individuals asymptomatically colonized with an ESBL-producing *E. coli* (ESBL^+^) and individuals who are not (ESBL^−^), determined by culture-based and molecular approaches. To avoid confounding factors from affecting study results, we selected samples from a large Dutch cross-sectional population study (PIENTER-3) for which 2751 fecal samples were used to culture ESBL-producing bacteria^10^. With this high number of samples available, we could apply stringent sample selection with regard to known confounders in microbiome studies such as antibiotic use, proton-pump inhibitor use, a variety of diets etc. Subsequently, we performed case control matching based on a variety of epidemiological and health related variables. We performed extensive functional and taxonomic profiling of the gut microbiome through metagenomics and metabolomics to investigate whether there are differences in the gut microbiome between matched ESBL^+^ and ESBL^−^ individuals.

## MATERIALS AND METHODS

### Sample collection

Samples were selected from a large Dutch population-wide study (PIENTER-3)^10^. This cross-sectional population study was carried out in 2016/2017, primarily designed to obtain insight into age-specific seroprevalence of vaccine-preventable infectious diseases. Out of the 98 included samples for the current study, 95 were stored in the freezer within 15 minutes after defecation, one person did not provide information on this and two individuals took longer than one hour to store their sample in the freezer. Samples were kept on average for 2.97 days (±2.82) (six individuals did not indicate this information) in people’s freezer before being delivered (on cold packs) to the mobile study team^10^. Fecal samples were kept on dry ice during transport to the National Institute for Public Health and the Environment and stored at −80°C the next day.

### Detection of ESBL-producing *Enterobacterales*

Details of the microbiological methods have been described elsewhere (Willems RPJ, van Dijk K, Dierikx CM, Twisk JWR, van der Klis FRM, de Greeff SC, Vandenbroucke-Grauls CMJE. Gastric acid suppression, lifestyle factors and intestinal carriage of ESBL and carbapenemase-producing Enterobacterales: a nationwide population-based study [Submitted]). Briefly, stool specimens were enriched by tryptic soy broth with ampicillin (50 mg/L) and then cultured on selective agar plates (EbSA, Cepheid Benelux, Apeldoorn). Next, up to five oxidase-negative morphotypes were subcultured, identified to species level, and tested for antimicrobial susceptibility using standard procedures (VITEK 2 system, bioMérieux, Marcy-L’Étoile, France). Antimicrobial susceptibility was classified according to European Committee on Antimicrobial Susceptibility Testing clinical breakpoints^11^. ESBL production was screened for with combination disk diffusion and confirmed by polymerase chain reaction (PCR); PCR was performed for the *bla*CTX-M, *bla*SHV and *bla*TEM groups^12^. ESBL testing was done according to the European Committee on Antimicrobial Susceptibility Testing guidelines^13^.

### Sample selection

2751 fecal samples were cultured for ESBL- or CPE-producing bacteria, of which 198 samples were positive. For the purpose of our study, we selected samples positive for ESBL-producing *E. coli*, resulting in 176 potential samples. Next, we applied stringent exclusion criteria for all samples based on variables known to affect the gut microbiome. Individuals were excluded based on the following criteria: current proton-pump inhibitor use, antibiotic use in the last three months, diarrheal symptoms in the last month (defined as at least three thin stools within 24 hours), vomiting in the last month, blood in stool during the last month, abdominal pain or nausea during the last month, use of any pre- or probiotics, consumption of a special diet (vegetarian, cow’s milk free diet, hen’s egg protein-free diet, gluten free, nut and/or peanut-free, lactose limited diet, diabetes-related diet, limited protein diet, limited fat and/or cholesterol diet, enrichment of dietary fiber, caloric restriction, low in sodium, easily digestible, coloring agent-free, enriched in energy/protein, ‘other diet’) and whether stool was stored in the freezer after defecation (samples were excluded if not stored in the freezer). This selection resulted in 51 ESBL^+^ samples for inclusion, which were subsequently matched to 51 ESBL^−^ samples using the R MatchIt package (v3.0.2) with the “nearest” method in the *matchit* function. Subjects were matched based on age, sex, having been abroad during the last 6 months (yes/no) and ethnicity. ESBL^−^ negative samples were selected using the same exclusion criteria. Three samples (1 ESBL^−^ sample and 2 ESBL^+^ samples) were further excluded as insufficient DNA was available for sequencing. One additional sample (ESBL^−^) was excluded as we discovered afterwards that this individual had provided ambiguous answers regarding dietary habits. The final dataset for analysis contained 49 individuals in each group.

### DNA extraction for metagenomic shotgun sequencing

DNA was extracted by mechanical disruption (repeated bead-beating) and purified in a Maxwell RSC instrument (Promega Benelux BV, Leiden, The Netherlands). The Maxwell RSC Blood DNA extraction kit was according to manufacturer’s instructions with several modifications, as follows. Fecal samples were thawed on ice and approximately 250 mg of well-homogenized fecal material was resuspended in S.T.A.R (stool transport and recovery buffer) buffer (Roche Diagnostics, Almere, The Netherlands), with 0.1 mm zirconia/silica beads and 2.5 mm glass beads. The fecal suspension was mechanically disrupted three times for one minute in a FastPrep-24 Instrument at room temperature and 5.5 oscillations, and maintained on ice after every cycle. Samples were further heated at 95°C for 15 minutes shaking at 300 rpm, and centrifuged for 5 minutes at full speed. Resulting supernatants (fecal lysates) were collected and the pellet was further resuspended in an additional 350 μl of S.T.A.R. buffer following the same procedure. Pooled fecal lysates were then transferred to the Maxwell RSC Instrument for further purification steps. Eluted sample was cleaned-up using the OneStep PCR Inhibitor Removal Kit (Zymo Research, Irvine, California), and DNA was quantified using a Quantus Fluorometer (Promega Corporation, Madison, WI, USA). Every extraction round included two negative DNA extraction controls (blank samples with S.T.A.R. buffer without any added fecal material) and two microbial mock communities as positive controls (ZymoBiomics Microbial Community Standards; Zymo Research, Irvine, California, USA).

### Metagenomic shotgun sequencing

Shotgun metagenomic sequencing was performed by GenomeScan B.V. (Leiden, The Netherlands) using the NEBNext® Ultra™ II FS DNA Library Prep Kit (New England Biolabs, Ipswich, Massachusetts, USA) and the NextSeq 500 platform (paired-end, 150bp). Two positive sequencing controls (ZymoBiomics Microbial Community DNA Standards; Zymo Research, Irvine, California, USA) and two negative sequencing controls (sterile water) were included. Average number of raw reads (of 98 samples and four positive controls) is 4,747,908 (range 2,565,232 – 62,035,096) and a median of 4,142,237 paired-end reads. Raw shotgun sequencing reads were quality checked using the FastQC (v0.11.9) and MultiQC (v1.8) tools, both before and after cleaning files for low-quality reads and human reads using the kneaddata (v0.7.10) tool with default parameters.

Taxonomic and functional annotation were performed on cleaned reads using the NGLess language (v1.2.0), associated tools and the Integrated Gene Catalog (IGC) database^14−18^. For taxonomic analysis, mOTUs (v2.5.1) was used with default parameters and unclassified reads (−1 category in mOTUs) were not included for downstream analyses^19^. Functional annotation was performed by aligning cleaned reads to the annotated IGC database (we annotated the IGC through eggNOG mapper v2.1.0 using default parameters and the “-m diamond” argument) using Burrows-Wheeler-Aligner MEM (BWA, v0.7.17)^17, 18, 20^. Unclassified reads were not taken into account for downstream analyses. Default parameters were used, apart from the ‘normalization’ argument, which was specified as normalization=“scaled”, which corrects for size of the feature (gene). Aligned reads were then aggregated using the Kyoto Encyclopedia of Genes and Genomes (KEGG), KEGG Orthology (KO) groups and Carbohydrate-active enzymes (CAZymes) annotations present in the IGC (features=“KEGG_ko” or features= “CAZy” argument in NGLess)^21, 22^.

Multi-locus sequence typing on *E. coli* was performed using the MetaMLST tool (default parameters). MetaMLST aligns sequencing reads against a database (which can be customized) of housekeeping genes to identify sequence types present in metagenomes. A custom *E. coli* database (Achtman MLST scheme) was created with MLST data from October 16^th^ 2020 (https://pubmlst.org/bigsdb?db=pubmlst_ecoli_achtman_seqdef)^23^. No sequence types could be reliably detected in the samples, likely due to the very low relative abundance of *E. coli* and the corresponding low number of reads and coverage of *E. coli*.

#### Resistome profiling

To profile the antimicrobial resistance genes in the metagenomes, cleaned reads were aligned to the MEGARes database (v2.00) using BWA MEM with default settings^17^. The resulting SAM file was parsed using the ResistomeAnalyzer tool (https://github.com/cdeanj/resistomeanalyzer) and the default threshold of 80% was used, meaning an antibiotic-resistance determinant was only included if at least 80% of the gene is detected in a sample^24^. Read counts originating from alignments to housekeeping genes associated with antimicrobial resistance (AMR) (e.g. *rpoB* and *gyrA*) that require single nucleotide polymorphisms to confer resistance were filtered out of the count table before downstream analyses, as previously reported^25^. Gene level data (e.g. *tetO*, *tetQ* and *tetW*) were used for calculating alpha and beta diversity metrics and for differential abundance analysis. For visualization purposes, gene level outputs were aggregated at the mechanism level (e.g. beta-lactams, mupirocin).

### Positive and negative controls for metagenomic sequencing

Eight mOTUs were detected in all four positive controls, exactly matching theoretical expectations. With regard to expected relative abundances, sequencing controls were, as expected, more accurate (average fold error of 1.14) than the DNA extraction controls (average fold error of 1.42 with underrepresentation of Gram-positive bacteria). The four included negative controls (two extraction controls and two sequencing controls) did not generate any reads. These results indicate good performance of sequencing, DNA extraction procedures and bioinformatic processing of the data.

### Metabolomics

The method for NMR analysis of fecal samples was adapted from the protocol developed by Kim et al. with a few minor adaptations^26^.

#### Sample preparation

Each feces-containing sample tube was weighed before sample preparation. To each sample tube 50 μl of 0.5 mm zirconium oxide beads (Next Advance, Inc.) and 750 μl of milli-Q water were added. Then, the tubes were subjected to bead beating for four sessions of one minute. The tubes were subsequently centrifuged at 18,000 g at 4°C for 15 minutes. For most samples, 600 μl of supernatant was transferred to new 1.5 ml Eppendorf tubes. In some cases the volume of available supernatant was slightly less. These tubes were centrifuged at 18,000 g at 4 °C for 1 hour. 270 μl of supernatant was added to 30 μl of pH 7.4 phosphate buffer (1.5 M) in 100% D_2_O containing 4 mM TSP-d_4_ and 2 mM NaN_3_. A customized Gilson 215 liquid handler was used to transfer the samples to a 3.0 mm Bruker NMR tube rack. The original sample tubes were cleaned, dried and weighed again.

#### NMR measurements

^1^H NMR data were collected using a Bruker 600 MHz Avance Neo/IVDr spectrometer equipped with a 5 mm TCI cryogenic probe head and a z-gradient system. A Bruker SampleJet sample changer was used for sample insertion and removal. All experiments were recorded at 300 K. A standard sample 99.8% methanol-d4 was used for temperature calibration before each batch of measurements^27^. One-dimensional (1D) ^1^H NMR spectra were recorded using the first increment of a NOESY pulse sequence^28^ with presaturation (γB1 = 50 Hz) during a relaxation delay of four seconds and a mixing time of 10 ms for efficient water suppression^29^. Initial shimming was performed using the TopShim tool on a random mix of urine samples from the study, and subsequently the axial shims were optimized automatically before every measurement. Duration of 90° pulses were automatically calibrated for each individual sample using a homonuclear-gated mutation experiment^30^ on the locked and shimmed samples after automatic tuning and matching of the probe head. 16 scans of 65,536 points covering 12,335 Hz were recorded. J-resolved spectra (JRES) were recorded with a relaxation delay of 2 s and 2 scans for each increment in the indirect dimension. A data matrix of 40 × 12,288 data points was collected covering a sweep width of 78 × 10,000 Hz. Further processing of the raw time-domain data was carried out in the KIMBLE environment^31^. The Free Induction Decay of the 1D experiment was zero-filled to 65,536 complex points prior to Fourier transformation. An exponential window function was applied with a line-broadening factor of 1.0 Hz. The spectra were automatically phase and baseline corrected and automatically referenced to the internal standard (TSP = 0.0 ppm). A sine-shaped window function was applied and the data was zero-filled to 256 × 16,384 complex data points prior to Fourier transformation. In order to remove the skew, the resulting data matrix was tilted along the rows by shifting each row (k) by 0.4992× (128-k) points and symmetrized about the central horizontal lines.

#### Metabolite quantification

Metabolites were quantified using KIMBLE and the results were checked by quantifying the same metabolites both in the JRES and in the NOESY1D experiments and in 10 randomly chosen spectra using the Chenomx NMR Suite version 8.6 (Chenomx Inc., Edmonton AB, Canada).

### Statistical analysis

#### Statistical software used for downstream analysis

Analyses and visualizations were performed in R (v4.0.4), using the following packages: phyloseq (v1.34.0), microbiome (v1.12.0), vegan (v2.5-7), tidyverse packages (v1.3.0), SIAMCAT (v1.10.0), table1 (v1.2.1) and ropls (v1.22.0)^32–38^. All analytical R code will be made publicly available upon acceptance of the manuscript. For all used tools, default parameters were used unless stated otherwise.

**Table 1:** Characteristics of participants included in the study. P-values were obtained using an independent t-test (for numerical variables) or Fisher’s exact test (for categorical variables).

|  | ESBL negative<br>(N=49) | ESBL positive<br>(N=49) | P-value |
| --- | --- | --- | --- |
| **Age (years)** |  |  |  |
| Mean (SD) | 44.1 (15.2) | 46.6 (15.3) | 0.43 |
| Median [Min, Max] | 45.0 [20.0, 74.0] | 46.0 [21.0, 74.0] |  |
| **Sex** |  |  |  |
| Male | 26 (53.1%) | 23 (46.9%) | 0.69 |
| Female | 23 (46.9%) | 26 (53.1%) |  |
| **Abroad in last 6 months** |  |  |  |
| Yes | 39 (79.6%) | 37 (75.5%) | 0.81 |
| No | 10 (20.4%) | 12 (24.5%) |  |
| **Ethnicity** |  |  |  |
| Dutch | 38 (77.6%) | 36 (73.5%) | 0.79 |
| First generation other-Western | 1 (2.0%) | 0 (0%) |  |
| Second generation other-Western | 2 (4.1%) | 3 (6.1%) |  |
| First generation Suriname+Aruba+Dutch Antilles | 3 (6.1%) | 3 (6.1%) |  |
| Second generation Suriname+Aruba+Dutch Antille: | 1 (2.0%) | 0 (0%) |  |
| First generation other non-Western | 4 (8.2%) | 7 (14.3%) |  |
| **Antibiotic use in the prior 3 to 12 months** |  |  |  |
| Yes | 6 (12.2%) | 7 (14.3%) | 0.77 |
| No | 43 (87.8%) | 41 (83.7%) |  |
| Do not know | 0 (0%) | 1 (2.0%) |  |

#### Community composition analysis of metagenomic data

We tested for differences in overall microbiota composition with permutational multivariate analysis of variance (PERMANOVA) using Bray-Curtis dissimilarity. As violation of the assumption of homogenous dispersions can lead to wrong conclusions regarding PERMANOVA, we first tested this assumption using the *betadisper* function of the vegan package. No heteroscedasticity was observed between the ESBL^+^ and ESBL^−^ group. To investigate both linear and non-linear patterns in the data, we performed dimension reduction using both principal coordinates analysis (PCoA) and t-distributed stochastic neighbor embedding (t-SNE), both based on Bray-Curtis dissimilarity. Alpha diversity indices were compared using independent t-tests.

#### Differential abundance analysis in metagenomic data

Differential abundance analysis of mOTUs, KO groups, CAZymes and resistance genes between ESBL^+^ and ESBL^−^ samples was performed using SIAMCAT on relative abundance matrices. Features (mOTUs, KO groups or CAZymes) had to be present in at least 25% of samples to be included in the analysis. Regarding resistome analyses, a gene had to be present in 10% of samples to be included, as the 25% prevalence cut-off was too stringent resulting in only fourteen genes included in the analysis. To correct for false discovery rate, p-values were corrected in all tests using the Benjamini-Hochberg procedure^39^.

#### Machine learning classifier on metagenomic data

We used obtained taxonomic and functional profiles for feature selection and construction of prediction models. To this end, least absolute shrinkage and selection operator (LASSO) logistic regression using the SIAMCAT package was performed to select predictive features and remove uninformative features based on species composition or functional profiles. Preprocessing was done by filtering mOTUs, KO groups, or CAZyme families which were present in at least 25% of samples. The vignette from SIAMCAT (https://siamcat.embl.de/articles/SIAMCAT_vignette.html) was followed^37^. In short, we performed data normalization using the “log.unit” method, 5-fold cross validation to split the data in several combinations of training and test data, trained the model using LASSO logistic regression (“lasso” parameter) and, lastly, made the predictions.

#### Metabolomics data

Metabolomic concentrations were first log10 normalized to reduce heteroscedasticity. Metabolite concentrations were subsequently centered and scaled to a mean of 0 and standard deviation of 1, as previously described^40^. Differences in concentrations between ESBL groups were tested using t-tests where p-values were corrected for multiple testing using two methods (to establish robustness of potential findings), namely Benjamini-Hochberg and Holm correction (with Holm correction being more conservative)^39, 41^. Next, we performed multivariate analyses using PCA and Partial Least-Squares Discriminant Analysis (PLS-DA). Lastly, random forest was applied to investigate whether ESBL^+^ and ESBL^−^ individuals could be accurately classified based on their respective metabolite profiles. As input to the random forest, normalized metabolite concentrations were used and, similarly as with metagenomic data, 5-fold cross validation was implemented in SIAMCAT.

## RESULTS

### Participant and ESBL-producing *E. coli* isolates characteristics

The original sample selection contained 51 individuals in each group, but three samples were not suitable for metagenomic sequencing due to too low DNA concentrations after extraction. One more individual had to be excluded due to ambiguous answers regarding dietary habits. Ultimately, this resulted in metagenomics data from 49 individuals per group. Demographic and participant characteristics were highly similar between the ESBL^+^ and ESBL^−^group and antibiotic use between the preceding three to twelve months was also evenly matched (Table 1). With regard to the ESBL-producing *E. coli* isolates that colonized our 49 ESBL^+^ participants, 44 carried a CTX-M-type. The majority of these were CTX-M-1 (25) and CTX-M-9 (18) and one could not definitively be typed (CTX-M-1 or CTX-M-8). Isolates of four individuals were negative for CTX-M genes and for one participant it could not be determined. Additional information on antimicrobial susceptibility of the strains can be found in Supplementary Table 1.

### No differences between the ESBL^+^ and ESBL^−^ individuals in bacterial species composition or diversity parameters

We investigated potential differences in microbiota composition and diversity between ESBL^+^ and ESBL^−^ samples. A total of 1178 species (mOTUs) were detected in our cohort. Overall bacterial composition at the family and genus level are shown in Figure S1. The most abundant species and their average relative abundance in this cohort were *Bifidobacterium adolescentis* (4.6% ± 6.9%), *Ruminococcus bromii* (3.4% ± 4.8%) undefined *Ruminococcaceae spp.* (2.9% ± 3.2%), *Eubacterium rectale* (2.7% ± 2.8%) and *Prevotella copri* (2.5% ± 5.7%). We did not observe differences in alpha diversity (observed mOTUs and Shannon index, Figure 1A and B), nor in beta diversity (PCoA and t-SNE, Figure 1C and D).

**Figure 1:**
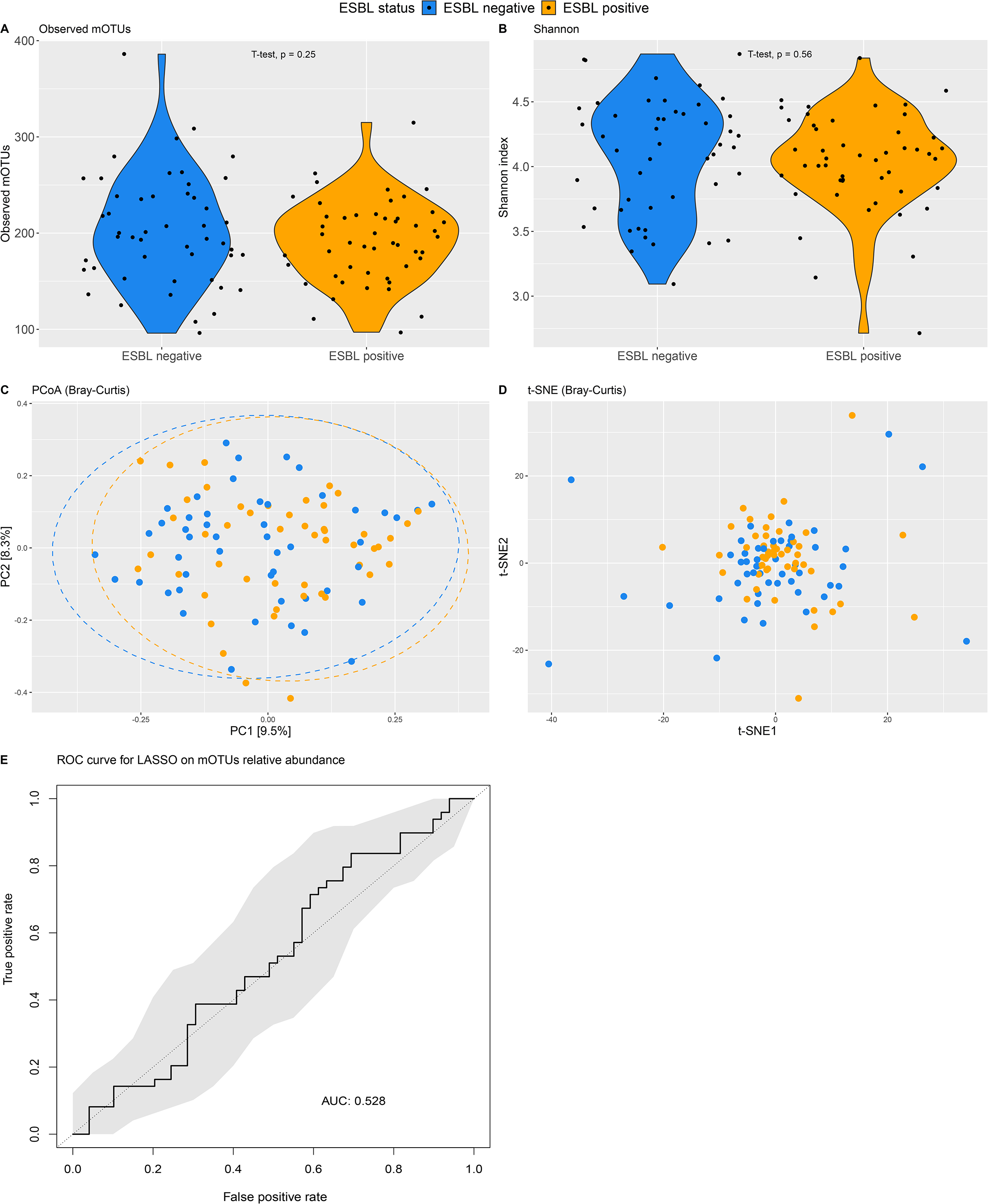
Taxonomic analyses between ESBL groups with comparisons of observed mOTUs (A) and Shannon index (B), unsupervised clustering using PCoA (C) and t-SNE (D) based on Bray-Curtis dissimilarity and the ROC curve for LASSO (E). The ROC curve shows the mean AUC value and its respective 95% CI.

Next, we investigated whether there were differences in relative abundance between the study groups at the species level (mOTUs). Prior to differential abundance testing, mOTUs were filtered based on a prevalence of at least 25%, resulting in 261 mOTUs (representing 22.2% of the total observed mOTUs). No significant differentially abundant mOTUs were detected (all corrected p-values > 0.7). In order to elucidate whether microbiota composition is predictive of ESBL carriage, a machine learning classifier (LASSO logistic regression) was applied to the filtered mOTUs relative abundance matrix, which provided an AUROC value of approximately random classification (AUROC of 0.53, Figure 1E), indicating that mOTUs relative abundance does not allow for reliable prediction of ESBL status.

### No differences in the resistome of individuals colonized by an ESBL-producing *E. Coli* and ESBL^−^ individuals

Of all cleaned reads, an average of 0.035% (±0.024%) reads per sample mapped against the MegaRes 2.0 database. There was no difference between ESBL groups in the average number of reads aligned to MegaRes 2.0 (independent t-test, p=0.84). A total of 98 unique antimicrobial resistance genes (ARGs) were detected with 17 different AMR mechanisms (e.g. beta-lactam), and the number of detected ARGs was not different between ESBL groups (independent t-test, p = 0.46) (Figure 2A). Overall ARGs profiles in the study groups assessed by plotting beta diversity, did not show a clear separation between ESBL groups (Figure 2B), which was confirmed by PERMANOVA (p=0.21). The most abundant ARGs and AMR mechanisms are visualized in Figure 2C and D. No differences in relative abundance of ARGs were found between the groups using differential abundance analysis (all corrected p-values > 0.4). Tetracycline resistance was most abundant in the resistomes (47.7% ± 24.7%, Figure 2C), followed by mupirocin resistance (33.7% ± 28.6%). Tetracycline resistance was conferred by several *tet* genes, while mupirocin resistance was conferred through the *ileS* gene. As it is known from literature that *Bifidobacterium* spp. can be intrinsically resistant to mupirocin through the *ileS* gene^42^, we analyzed the correlation between the relative abundance of *Bifidobacterium* (at genus level) and the *ileS* gene, which was indeed high (R=0.78, p<2.2×10^16^) (Figure S2). We then moved on to investigate functional profiles of our participants.

**Figure 2:**
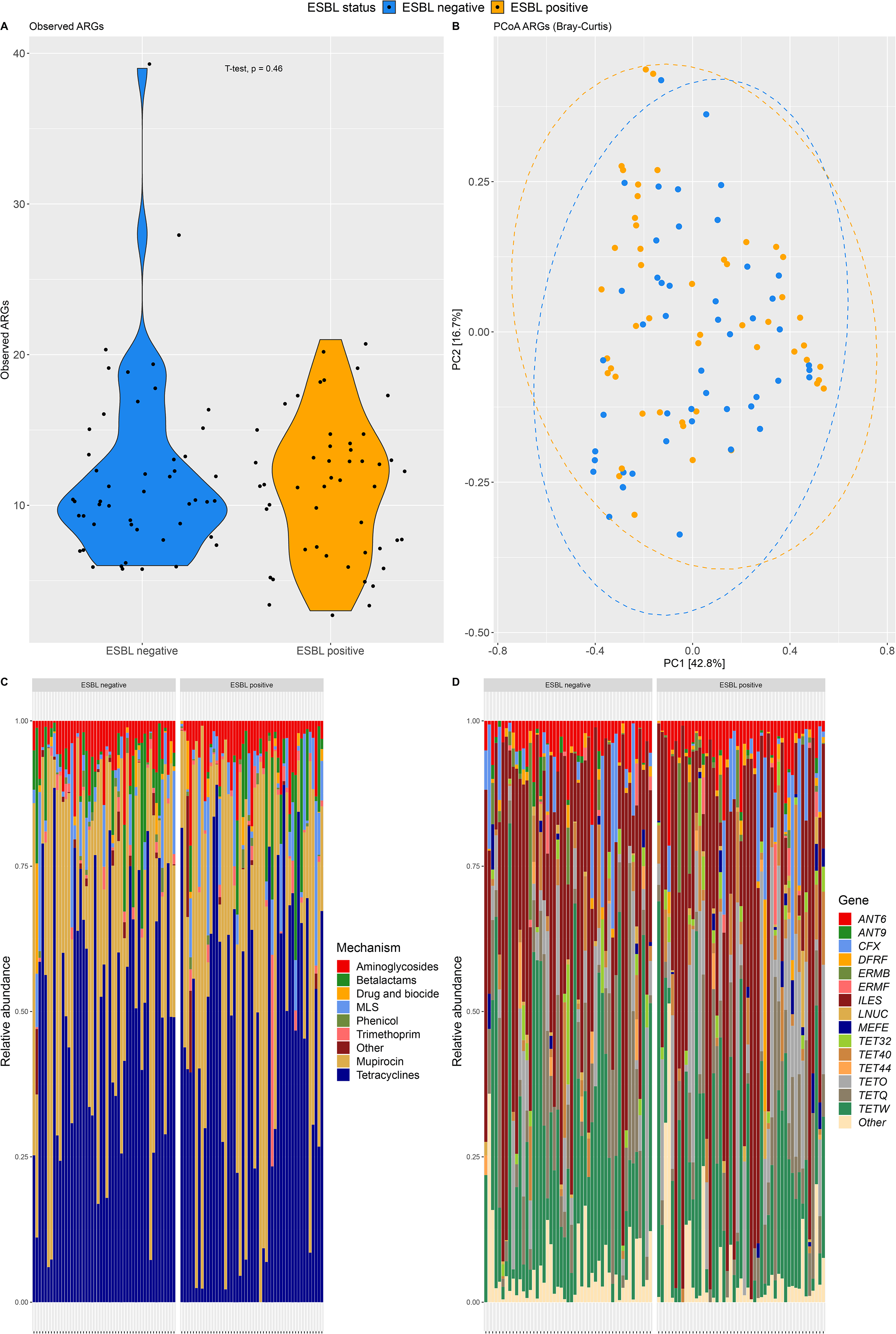
Resistome analyses with comparisons of the number of detected ARG (A), resistome diversity (B) and overviews of the most abundant resistance mechanisms (C) and resistance genes (D).

### No differences between the ESBL^+^ and ESBL^−^ individuals in functional capacity of the microbiome

To compare the functionality of the gut microbiome between the study groups, cleaned reads were mapped against the annotated IGC database. On average, 95.8% (±1.7%) of reads aligned against the IGC, and the aligned number of reads was not different between ESBL groups (independent t-test, p=0.23). From the aligned reads, 49.2% (±2.2%) aligned against a gene annotated by a functional group (KO group) and this was not different between ESBL groups (independent t-test, p=0.13). There was no difference in overall functional profiles between the groups (PERMANOVA, p=0.19). 8450 KO groups were detected and after filtering on 25% prevalence, 5179 KO groups remained for differential abundance testing. No KO groups were significantly differentially abundant between ESBL groups (all corrected p-values > 0.2). To identify functional groups predictive of ESBL status, LASSO logistic regression was applied to the relative abundance matrix of KO groups. No accurate prediction model could be constructed (AUROC of 0.61), indicating that the functional groups do not contain information allowing for prediction of ESBL status.

### No functional differences in Carbohydrate Active Enzymes (CAZymes) between the ESBL^+^ and ESBL^−^ group

From the aligned reads, 2.1% (±0.2%) aligned against a gene annotated to a CAZyme family and this was not different between ESBL groups (independent t-test, p=0.48). A total of 109 CAZyme families were detected with a mean of 77.7 (±5.7) per individual, with no differences between ESBL groups (independent t-test, p=0.34) (Figure 3A). The three most abundant CAZymes in our study were glycoside hydrolase (GH)13 (19.4% ± 3.3%), GH3 (11.4% ± 1.6%) and GH31 (6.2% ± 0.9%) (Figure 3C), corresponding to breakdown of starch and glycogen (GH13) and breakdown of plant cell wall glycans (GH3 and GH31)^43^. Variation in CAZyme relative abundance profiles could not be explained by ESBL group (PERMANOVA, p=0.57, Fig 3B). Compositional plots based on the top 20 most abundant CAZymes were highly similar between the ESBL groups (Figure 3C), and no differences in relative abundance of individual CAZyme families was observed (all corrected p-values > 0.6). To identify potential drivers of ESBL-producing *E. coli* colonization we used LASSO logistic regression on relative abundances CAZymes, which did not result in an accurate prediction model (AUROC of 0.56). This indicates there is only very low to no predictive power in relative abundances of CAZymes with regard to ESBL status.

**Figure 3:**
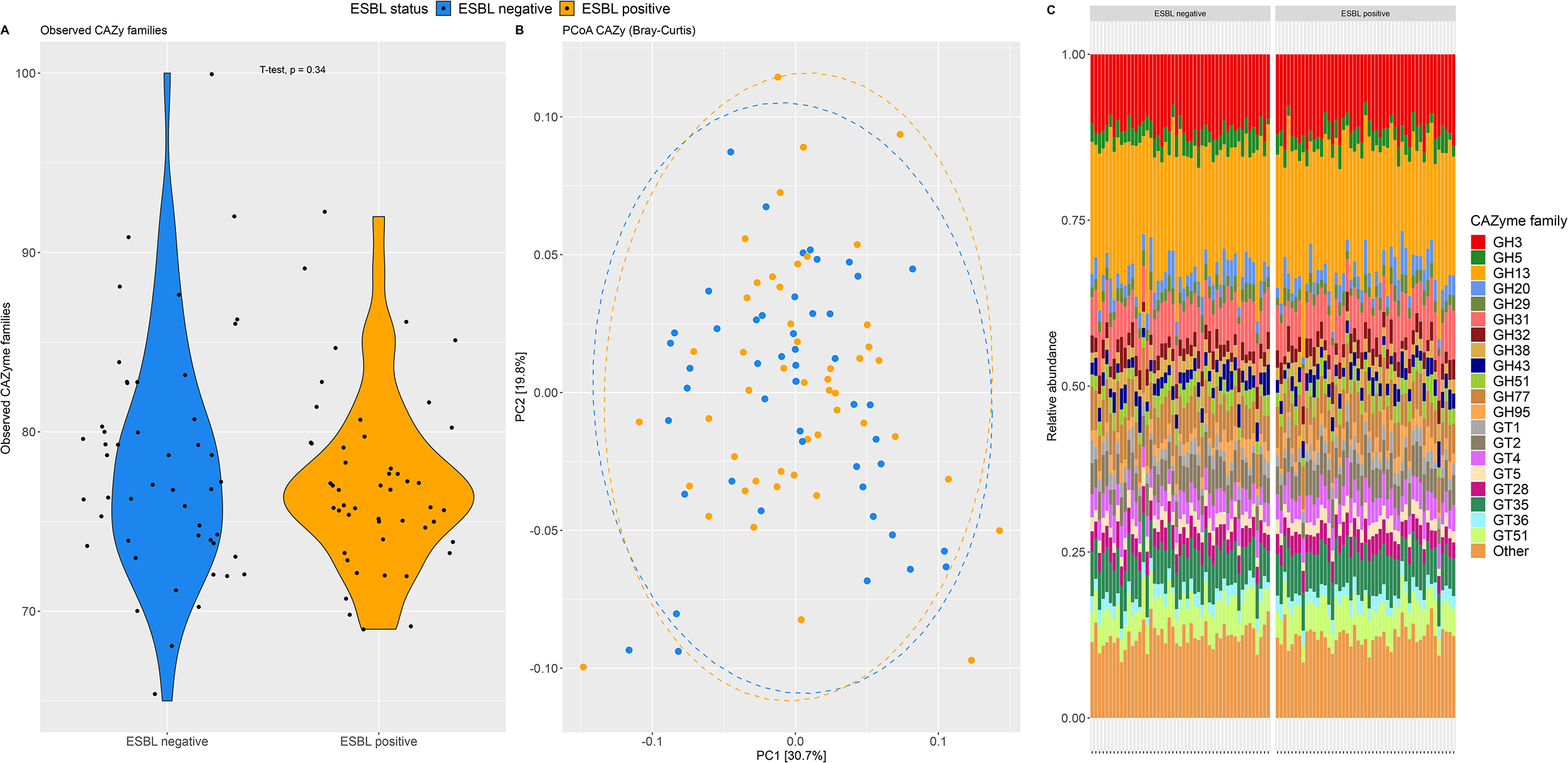
Overview of analyses based on CAZyme repertoire with a comparison of number of CAZyme families (A), PCoA based on Bray-Curtis dissimilarity (B) and a compositional plot to show the consistency of CAZyme families across participants. GH: Glycoside hydrolase, GT: glycosyl transferase.

### Metabolomics profiling shows no clear differences between ESBL groups at the functional level

For metabolomic analysis we quantified metabolite concentrations in all individuals, except for one ESBL^+^ sample that was excluded as a good quality NMR spectrum could not be recorded due to shimming problems. First, to investigate whether any differences in metabolite concentrations existed between ESBL groups, we performed univariate testing (independent t-tests). These results strongly depended on the method used for multi-error correction (11 metabolites were significantly different at p=0.048 with Benjamini-Hochberg, but none with Holm) (Figure S3 and Figure S4).

Unsupervised dimensionality reduction using PCA was performed to investigate whether any separation could be observed based on ESBL carriage (Fig 4A). Over 46% of the metabolome variation could be explain on the first principal component, with some separation of the study groups. However, supervised analysis using a PLS-DA indicates that no predictive value could be obtained for class separation based on two PLS components (Q2Y = −0.06). Lastly, we performed a random forest prediction model to investigate whether ESBL status could be predicted based on metabolite profiles, but this was not the case (AUROC = 0.61) (Figure 4B). Altogether, minor differences in metabolite concentrations could be detected using t-tests, but these were dependent on the method applied for correction for multiple testing. PCA between the ESBL groups showed a small overall signal, but no predictive value could be confirmed by both PLS-DA and random forest.

**Figure 4:**
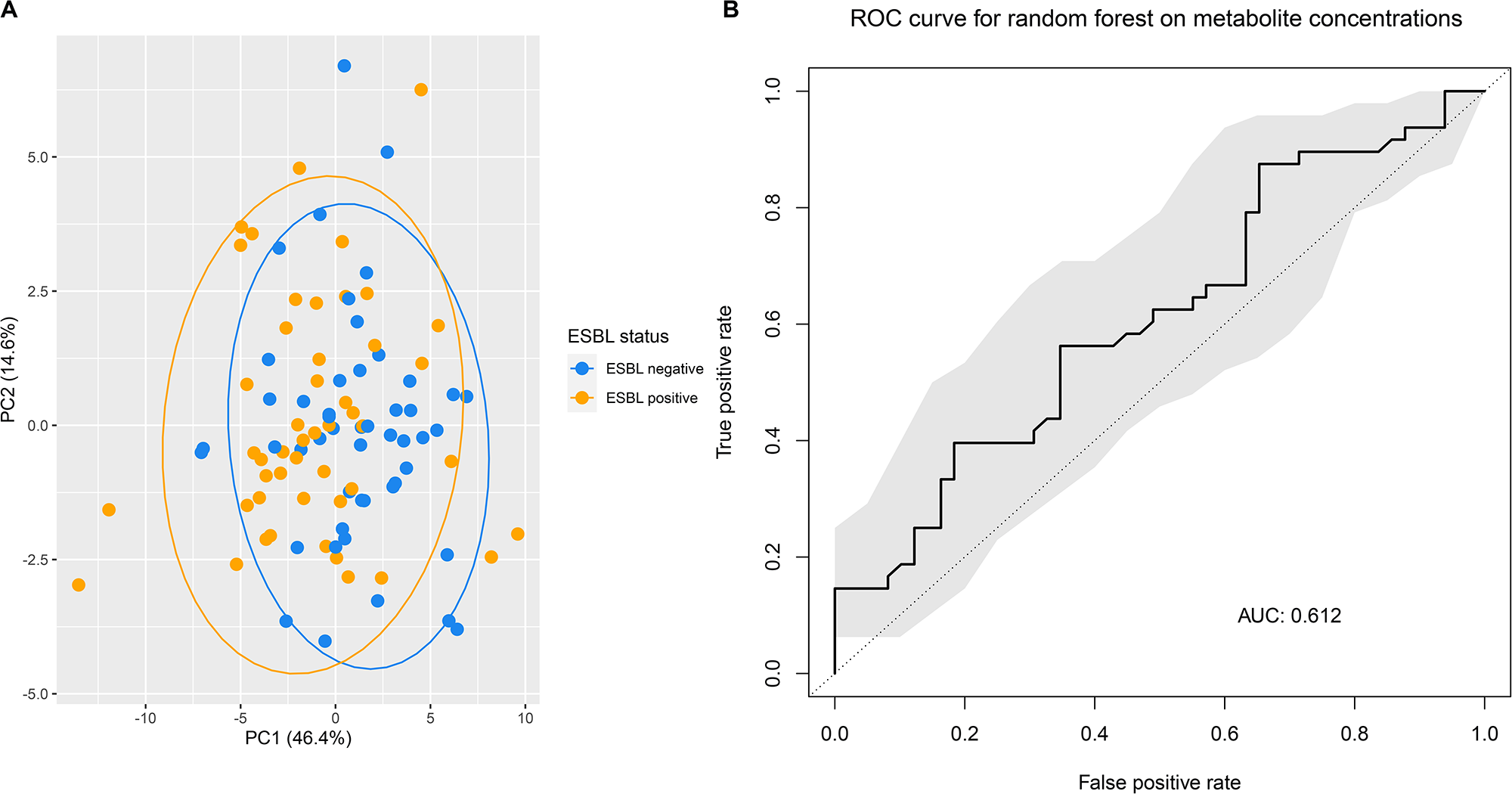
Metabolomic analyses with PCA (A) and the ROC curve of random forest based on metabolite profiles (B). The ROC curve shows the mean AUC value and its respective 95% CI.

## DISCUSSION

We present a unique study investigating differences in the gut microbiome and metabolome between individuals asymptomatically colonized by an ESBL-producing *E. coli* and matched non-colonized individuals. Importantly, in contrast to previous studies on this topic, we applied stringent inclusion criteria and matched ESBL^+^ individuals with ESBL^−^ individuals on important epidemiological variables, which minimized the chance for observing effects which could be attributed to confounding variables. The combination of metagenomics and metabolomics allowed for a deep molecular resolution of the gut microbiome, both at the taxonomic and functional level. We show that there is no difference in the gut microbiome of individuals asymptomatically colonized with an ESBL-producing *E. coli* as compared to individuals who are not colonized.

Confounding factors may, at least partially, be the reason for the previously reported differences in microbial signatures associated with protection from asymptomatic colonization by ESBL-producing bacteria and MDROs across different studies. It must be noted that these studies have mostly investigated vulnerable patient populations, such as nursing home residents and hospitalized patients. In such populations it is very complex to disentangle observed differences between colonized and non-colonized individuals from differences due to confounding variables (such as comorbidities and medication) between compared individuals^6, 8, 44—46^. In our study we excluded individuals based on many microbiome-influencing clinical factors, and performed matching on several clinical variables, as recently recommended for cross-sectional microbiome studies^9^. In this way, we could study the effect of colonization of ESBL-producing *E. coli* in isolation and convincingly show that no differences exist in the gut microbiome between colonized and non-colonized individuals.

In addition, previous research has generally not focused on species-specific colonization resistance, but rather on a broad category of MDROs (such as ESBL-producing *Enterobacterales*)^6, 8, 44–46^. Given the large genomic diversity within species^47^, let alone within the order of *Enterobacterales*, it is highly unlikely that a common mechanisms exists which could prevent colonization of e.g. both ESBL-producing *Klebsiella pneumoniae* and ESBL-producing *E. coli*. Therefore in the current study we focused on a single species (*E. coli*), rather than a broad group of ESBL-producing *Enterobacterales*.

Microbiome composition of individuals in our study population reflects that of other population cohorts in general. For example, *B. adolescentis* has been previously described in another Dutch cohort as the most abundant bacterial species, with an average relative abundance of 9.51% (±10.8%)^48^. In addition, *P. copri*, *R. bromii* and *E. rectale* were also highly abundant and prevalent, in line with the findings in the current study^48^.

The resistome profiles identified in our study also corresponded well with what is generally described in literature, with tetracyline resistance being the most abundant resistance mechanism in the human gut^49–51^. The observed high relative abundance to mupirocin in our study could be explained by the intrinsic resistance of *Bifidobacterium spp*. to this, of which relatively high abundances were observed in this cohort.

We show that despite inter-individual variation in taxonomic profiles, the functionality of the microbiome as assessed by the relative abundance of CAZyme families, is highly consistent between individuals. These finding are in line with previous findings showing functional similarity at the metabolic level despite taxonomic diversity^52, 53^.

This study is, to our knowledge, the first study to profile the gut metabolome in relation to colonization of ESBL-producing *E. coli*. We did not observe a relation between the metabolome, or any specific metabolite, and ESBL status. For other enteric pathogens, like *Salmonella enterica* serovar Typhimurium and *C. difficile*, specific metabolites have been shown to be strongly related to colonization resistance in rodent models^54, 55^. It should however be mentioned that these are infection models rather than asymptomatic colonization models, which would better represent our study.

A limitation of our study is that we do not have longitudinal data on the microbiome of these participants, and are therefore unable to make any statements about the duration of colonization of ESBL-producing *E. coli* and associations with the gut microbiome in time. This is particularly relevant considering the large variation in the duration of colonization between individuals^56, 57^. It could be speculated that individuals who are long-term colonized have a different gut microbiome than individuals who are only colonized for a short period of time, although there is no clear evidence for this in literature to our knowledge. Furthermore, longitudinal observations would allow us to identify changes occurring at the compositional and functional level when asymptomatic carriage turns into active infection or when people become decolonized. Lastly, one would have ideally have microbiome data of an individual shortly before an ESBL-producing *E. coli* would colonize and at time of colonization, so that microbiome changes within an individual can be investigated. Secondly, we do not have whole-genome sequencing data of the ESBL-producing *E. coli* isolates, which prevents us from placing these data into a broader epidemiological context. For example, if the majority of isolates would be sequence type (ST)131, an endemic ST, this would be valuable extra information and further extend the clinical relevance of our findings.

This study is however unique in the fact that ESBL^+^ and ESBL^−^ individuals were selected from a large Dutch cohort (n= 2751), and therefore we could apply stringent inclusion criteria and match the two groups on several demographic and clinical variables. To the best of our knowledge, this is one of very few studies in the microbiome field that applied such a stringent study setup. This setup ensured that the potential effect of confounding factors was minimized. In addition, this study is the first to investigate differences in the gut microbiome and metabolome between individuals colonized by an ESBL-producing *E. coli* and non-colonized individuals using a combined approach of metagenomics and metabolomics. Therefore, it provides insight into both the composition and function of the gut microbiome.

## CONCLUSIONS

Our study shows that there are no differences in the gut microbiome or metabolome of individuals who are, or are not, asymptomatically colonized by an ESBL-producing *E. coli*. We hypothesize that microbiome-mediated colonization resistance may therefore not be as relevant against ESBL-producing *E. coli* as it is for other enteric pathogens (like *C. difficile* and vancomycin-resistant *Enterococcus*), although longitudinal studies or controlled human colonization models are necessary to confirm this hypothesis.

## ACKNOWLEDGEMENTS

We gratefully acknowledge all individuals who participated in the PIENTER-3 study.

## AUTHOR CONTRIBUTIONS

QD, RZ and EK conceived and designed the study. RZ, SF and EK supervised the study. RW and CV performed detection of ESBL-producing bacteria and RW aided in sample selection. AV and MG performed metabolomics analysis and AV aided in statistical analysis of metabolomics. FK and EF coordinated sample collection. JK performed DNA extraction and related laboratory procedures. QD performed sample selection, processed and analyzed metagenomic data, performed statistical analysis of metabolomics data, created figures and wrote the manuscript. SN aided in metagenomic data processing and analysis. QD, RZ, RW, AV, SN, FK, EF, JK, AV, MG, CV, SF and EK discussed results and implications. All authors contributed to and approved the manuscript.

## CODE AND DATA AVAILABILITY

All raw metagenomic data will be released under PRJEB44119 upon acceptance of the manuscript. All R code and necessary data files to reproduce the analyses and figures will be uploaded to GitHub upon acceptance of the manuscript.

---

**Table S1:** Additional information on antimicrobial susceptibility of ESBL-producing *E. coli* strains based on clinical breakpoints of EUCAST.

**Figure S1:** Compositional plots of taxonomy at family level (A) and genus level (B) for all participants in the current cohort, facetted by ESBL status. The 20 most abundant families and genera across all individuals are displayed. ‘Other’ indicates the sum of all bacterial families or genera not indicated in the respective plot.

**Figure S2:** Spearman correlation plot between relative abundance of the *ileS* gene and *Bifidobacterium* (at genus level). Correlation coefficient (R) and significance are indicated in the plot.

**Figure S3:** Metabolite concentrations (after log10 normalization and scaling) in μmol/g feces of all measured metabolites per ESBL group. T-tests with Benjamini-Hochberg adjustment for multi-error correction were performed to obtain indicated p-values.

**Figure S4:** Metabolite concentrations (after log10 normalization and scaling) of all measured metabolites per ESBL group. T-tests with Holm adjustment for multi-error correction were performed to obtain indicated p-values.

