## Supplementary figures and images for "Asymptomatic gut colonization by extended-spectrum beta-lactamase-producing *Escherichia coli* is not associated with an altered gut microbiome or metabolome in Dutch adults"

### Figure_S1

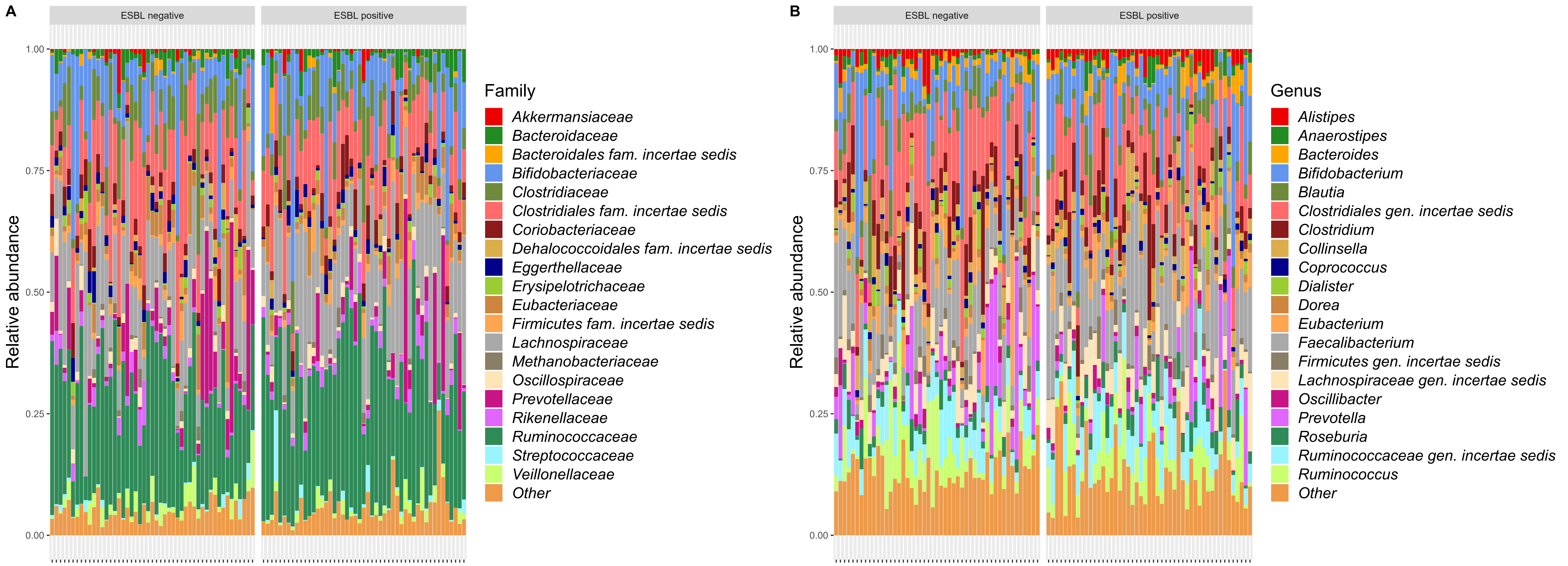

### Figure_S2

Correlation between relative abundance of *ileS* gene and *Bifidobacterium*

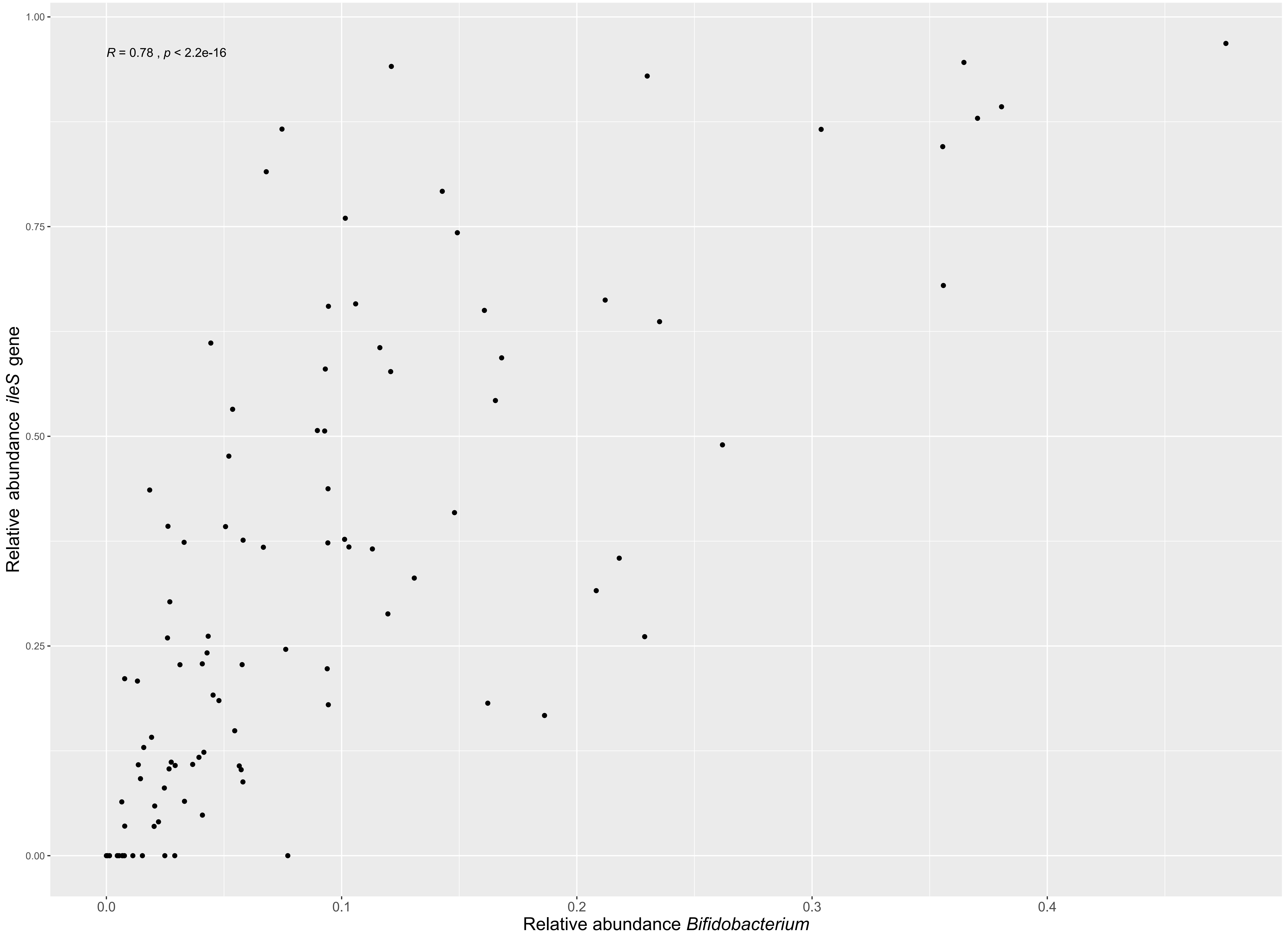

### Figure_S3

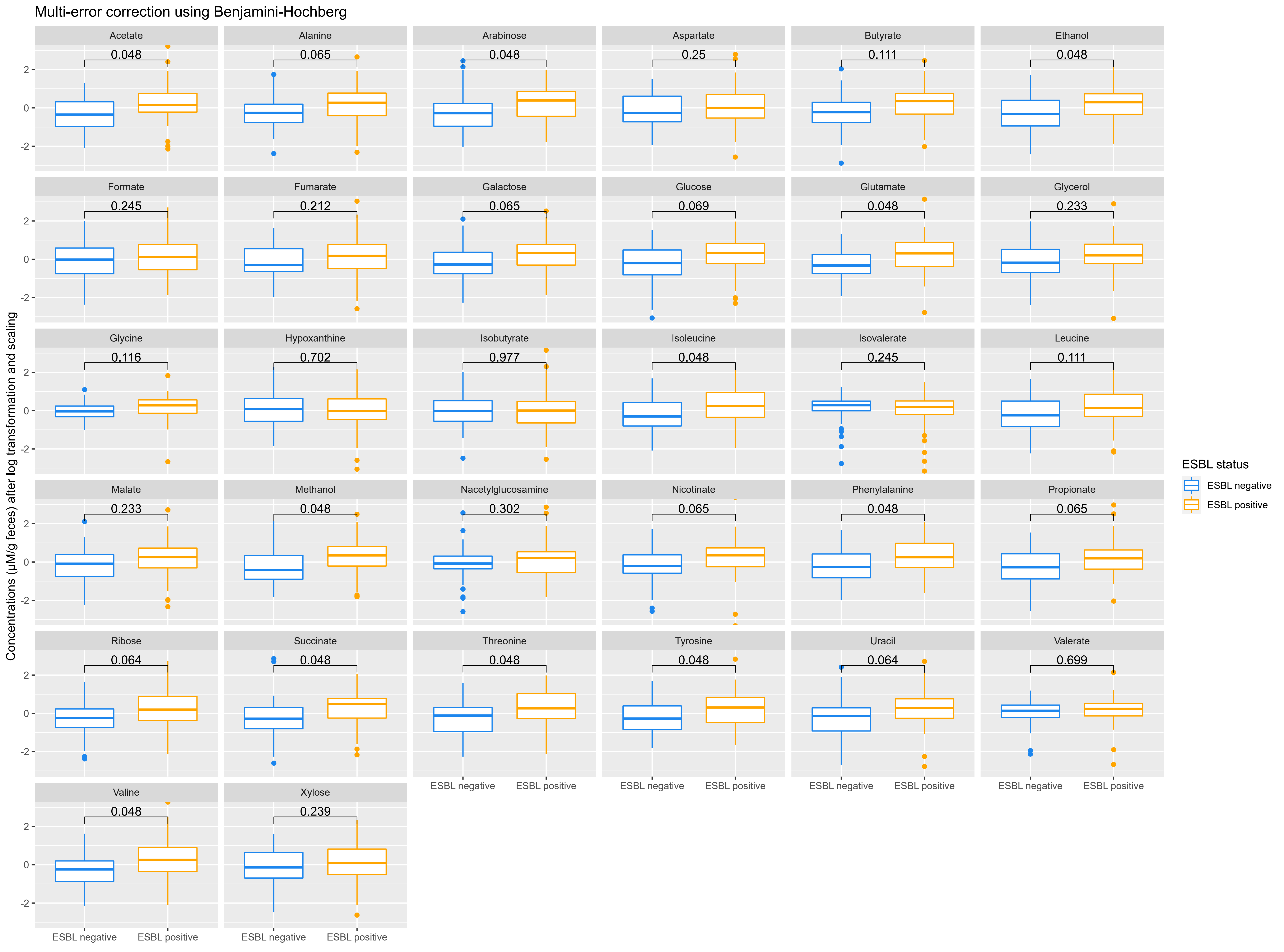

### Figure_S4

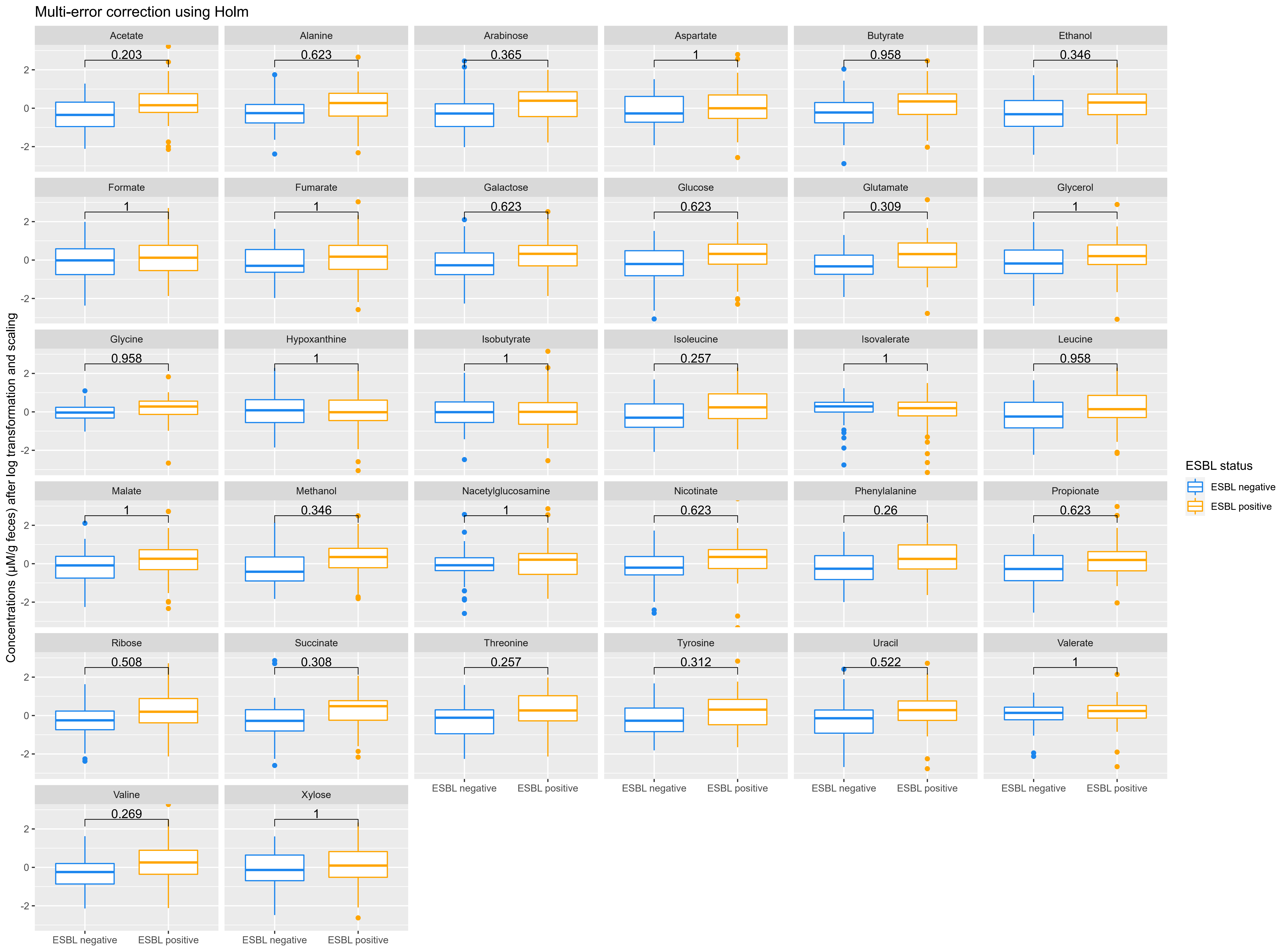
